# Armor reduction and pelvic girdle loss in a population of threespine stickleback *(Gasterosteus aculeatus*) from western Newfoundland, Canada

**DOI:** 10.1101/2022.09.20.508658

**Authors:** RJ Scott, GE Haines, CA Trask

**Affiliations:** School of Science and the Environment, Grenfell Campus, Memorial University of Newfoundland and Labrador, Corner Brook NL, Canada, A2H 6K8

## Abstract

We describe the antipredator armor of a unique population of threespine stickleback (*Gasterosteus aculeatus*) from Narrows Pond in western Newfoundland and compare traits for this population to nearby populations from marine and freshwater systems. After standardizing for length, Narrows Pond stickleback are shallower bodied and have shorter dorsal spines than stickleback from the other populations. Also, though the number of armor plates for Narrows Pond stickleback is greater than for typical low-plate morphs, the size of the lateral plates for Narrows Pond stickleback is much smaller. Finally, most (nearly 75% of sampled individuals) Narrows Pond stickleback do not have a pelvic structure (bilateral pelvic plate, ascending process, and ventral spine) and the remaining individuals have greatly reduced pelvic girdle whereas all individuals from the other populations possessed complete pelvic structures.

## Introduction

Freshwater fishes in temperate, post-glacial lakes are an excellent example of locally adaptive population divergence (Robinson and Wilson 1993; Foster et al. 1998; Robinson and Parsons 2002; Losos 2010; Arostegui and Quinn 2019). Repeated examples of adaptation to local environmental conditions provides an opportunity to understand the microevolutionary processes that produce this variation and ultimately could help understand the importance of those mechanisms for reproduction of new species. The repeated parallel variation exhibited by freshwater populations of threespine stickleback (*Gasteroteus aculeatus*) is particularly useful for understanding the roles of genetics, population history, gene flow and natural selection, which vary in importance among many systems (Foster et al. 1998, Losos 2010, Stuart et al. 2017, Fang et al. 2020).

Variation in the bony antipredator armor among freshwater populations of threespine stickleback has been particularly well studied. Individuals in marine populations, the ancestral form of all freshwater populations throughout the species’ range, possess a set of antipredator armor comprised of a set of lateral bony plates (30-34), a bony pelvic girdle, and dorsal and pelvic spines (Fig. 1). The pelvic girdle is comprised of two ventral bony plates fused along the fish’s midline and composed of anterior, posterior, and ascending processes (Bell 1987). The ascending processes often overlap 1-3 of the lateral plates, which in turn are typically partly overlapped by the basal plates of the dorsal spines, creating a rigid ring encircling the body (Reimchen 1983). The pelvic spines are anchored to the ventral portion of the pelvic girdle at the base of the ascending processes, and the dorsal spines are anchored to the dorsal pterigiophores and basal plates (Fig. 1). Freshwater populations vary in the expression of antipredator armor traits from those that are similar to the marine form, to those that have reduced number of armor plates (typically 5-8 and lacking a caudal keel), reduced size of the pelvic girdle, and smaller spines, and in rare instances loss of some or all of the pelvic girdle complex (e.g. Bell and Orti 1994; Klepaker et al. 2012).

**Figure 1.**
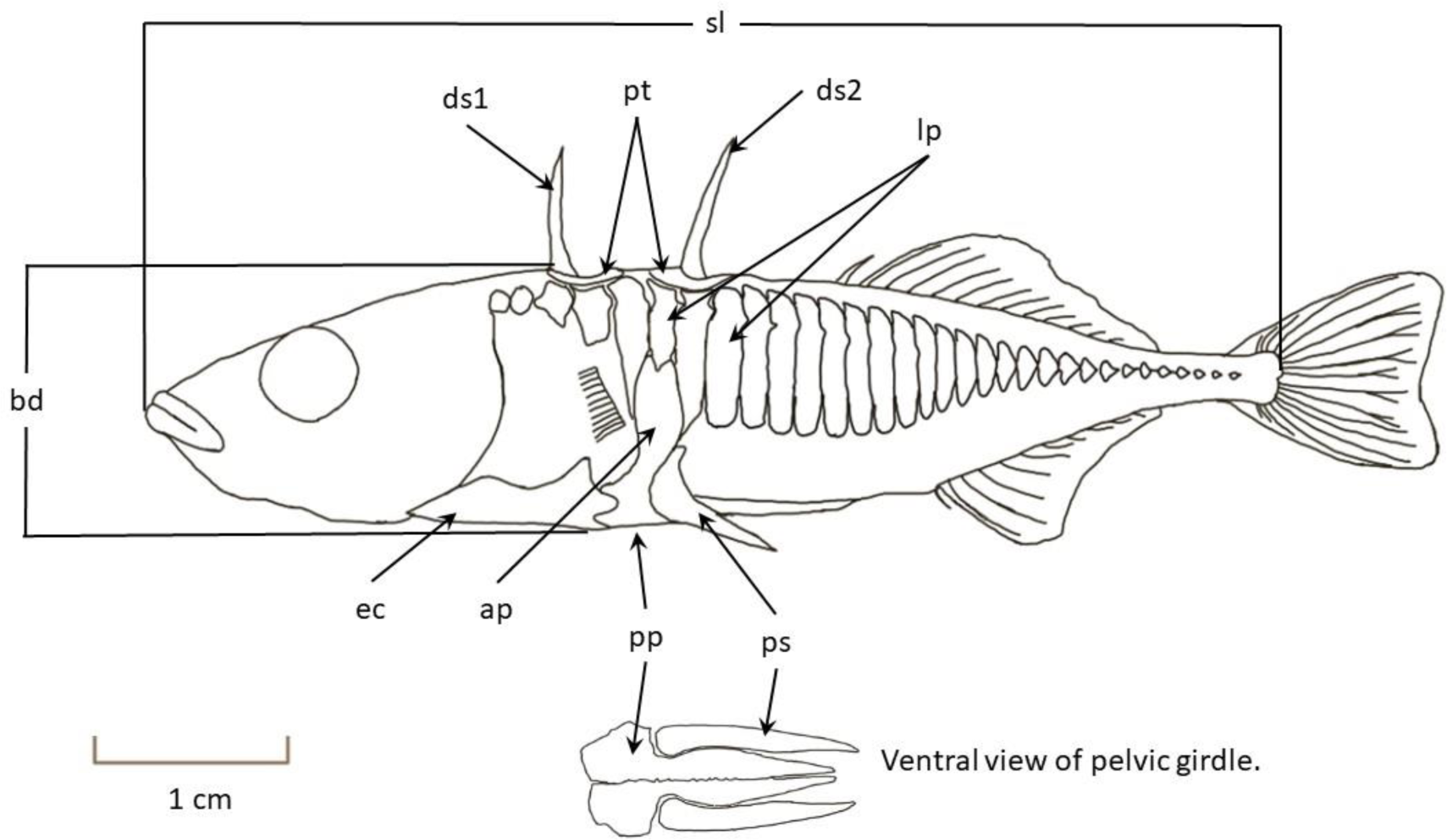
Line drawing of a typical full-plated threespine stickleback showing the suite of antipredator armor structures (ds1=1^st^ dorsal spine; ds2=2^nd^ dorsal spine; pt=dorsomedial pterigiophores; lp=lateral plate; ap=ascending process, pp=pelvic plate; ps=pelvic spine).

Pelvic girdle reduction and loss has been observed primarily in the eastern Pacific and northern European threespine stickleback, although it is rarely present elsewhere in other stickleback species (Klepaker 2013). Although pelvic reduction has been observed in eastern North America, it has been documented in only two nearby populations in Québec (Edge and Coad 1983). We found a population of threespine stickleback with pelvic girdle reduction during a larger survey examining variation in antipredator armor and body shape among stickleback from the west coast of Newfoundland (Fig. 2 for an example of stickleback from this population). Here, we compare antipredator armor in general, and pelvic girdle traits specifically, between the pelvic reduced population and other nearby freshwater and marine stickleback populations.

**Figure 2.**
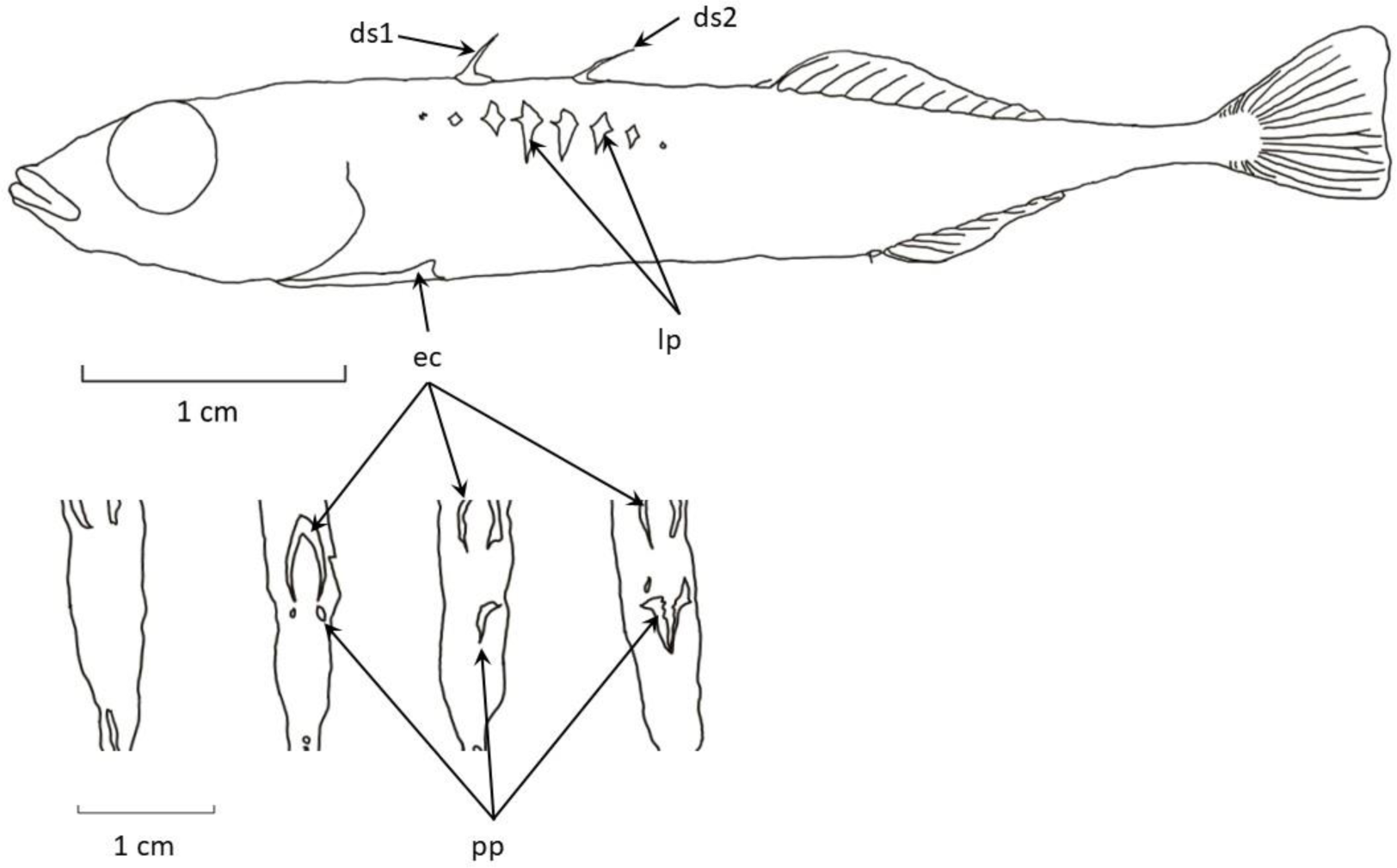
Line drawing of a Narrow Pond stickleback (top) showing the suite of antipredator armor structures (ds1=1^st^ dorsal spine; ds2=2^nd^ dorsal spine; lp=lateral plate; ec=ectocorracoid; pp=pelvic plate) and examples of pelvic girdle expression (bottom). Pelvic girdles are arranged in increasing expression from left to right (absent to maximum observed expression).

## Methods

We collected stickleback from Narrows Pond (the system containing the pelvic reduced populations) and from several nearby freshwater and marine locations during June and July between 2013 and 2020 (Fig. 3). The nearby freshwater populations sampled represented by complete (High Elevation Pond), partial (Rocky Harbour Pond), and low-plated (Bonne Bay Little Pond, Tilt Pond, Western Brook Pond) stickleback plate morphs (Barrett 2010). Fish were collected using un-baited standard Gee-type minnow traps (1/8” and 1/4” mesh) set for between 6 and 24 hours in nearshore locations 0.5 to 1.5 m deep. Fifty to 100 individuals were euthanized (over-dose of MS-222) and preserved in 10% formalin in the field. Later, we re-hydrated the samples in deionized water and stained them with Alizarin Red-S following methods outlined in Song and Parenti (1995). Between 15 and 25 individuals from each sample were photographed (Cannon 5D Mark III with a 100 mm macro lens or Nikon D600 with a 90 mm macro lens) and we used ImageJ (Schneider et al. 2012) to measure various traits (shown in Fig.s 1 and 2). We also visually assessed the combined pelvic score (Klepaker and Østbye 2008) of 100 individuals from each of Narrows Pond and four other populations representing a marine population and three freshwater populations, one with either a complete set, one with a partial set and one with a low set of armor plates.

**Figure 3.**
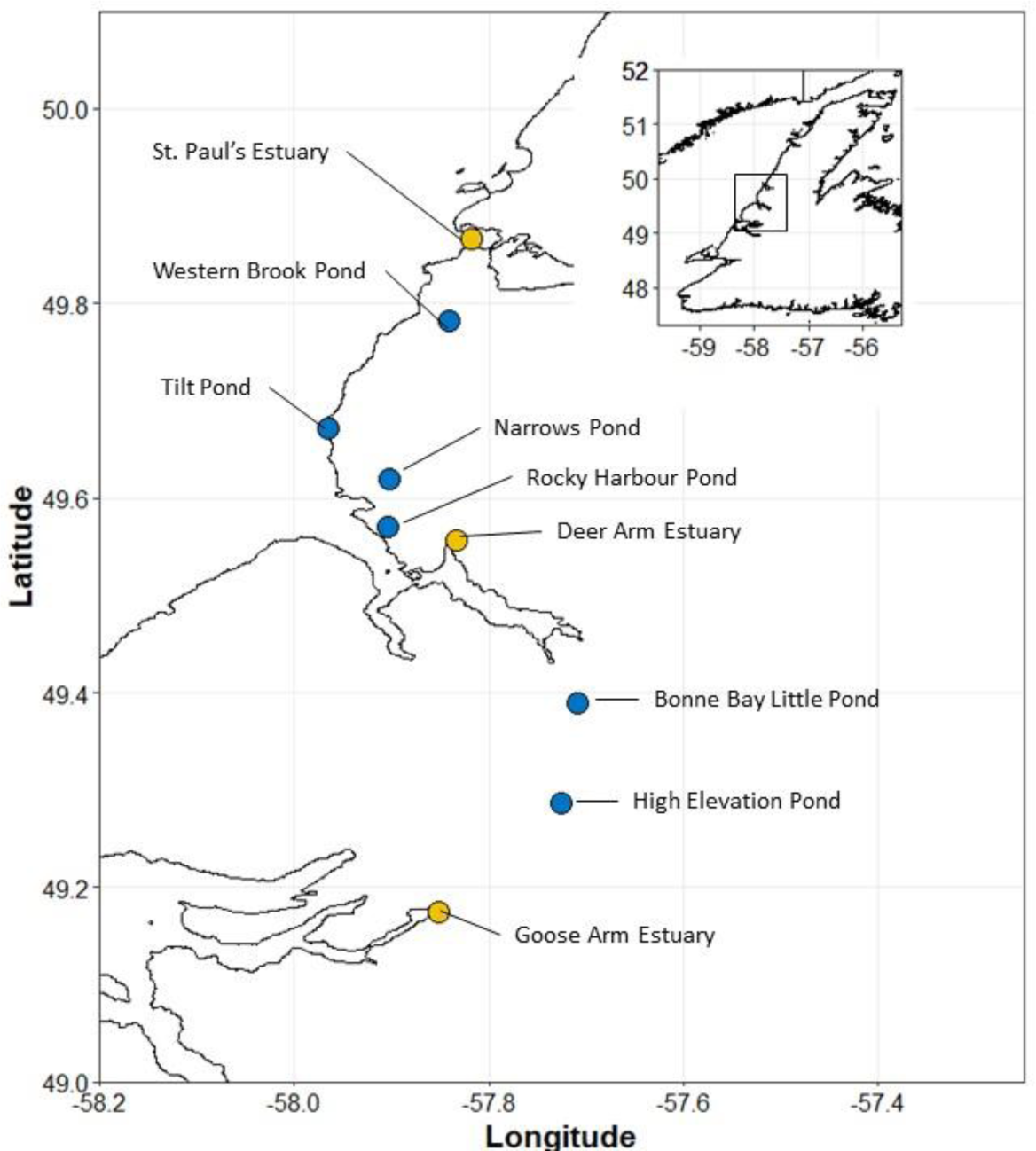
Map of western Newfoundland (inset) showing location of sampling sites in this study. Yellow symbols represent marine locations and blue symbols represent freshwater locations.

Statistical analyses were performed using the RStudio environment (version 2021.09.2+382; RStudio Team 2020) for R (version 4.1.3; R Core Team 2022). We adjusted for allometric effects following the method outlined by Lleonart et al. (2000) with modifications described in Stuart et al. (2017). Size adjusted values for each individual across all linear traits were calculated using the following:

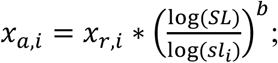

where *x*_*a,i*_ is the size adjusted trait value for individual *i, x*_*r,i*_ is the raw trait value for individual *i, SL* is the average standard length for all individuals measured, *sl*_*i*_ is the standard length for individual *i* and *b* is the slope of the relationship between the log of specific trait against the log of standard length. The slope of the relationship was determined using linear regression of each log transformed trait against the log transformed standard length across all sampled individuals. Size-adjusted traits (ln transformed) were compared among types using MANOVA (Quinn and Keough 2002).

## Results

The traits measured were all positively correlated with Pearson’s r ranging from 0.514 to 0.952 for raw trait values and from 0.393 to 0.949 for size-adjusted trait values (Table 1).

**Table 1.** Correlation (Pearson’s r) among the armor variables measured. Values below the diagonal are based on correlation among raw trait values and those above the diagonal are base on size-adjusted trait values.

|  | Pelvic<br>girdle width | Left<br>Pectoral<br>Spine | Pelvic<br>girdle<br>length | Ascending<br>process<br>length | Ascending<br>process<br>width | 2<sup>nd</sup> dorsal<br>spine | Mean<br>lateral plate<br>length | 1<sup>st</sup> dorsal<br>spine | Body depth | Standard<br>length |
| --- | --- | --- | --- | --- | --- | --- | --- | --- | --- | --- |
| Pelvic girdle width | 1.000 | 0.927 | 0.945 | 0.925 | 0.898 | 0.803 | 0.861 | 0.78 | 0.738 | 0.512 |
| Left Pectoral Spine | 0.938 | 1.000 | 0.949 | 0.94 | 0.881 | 0.874 | 0.812 | 0.862 | 0.633 | 0.4 |
| Pelvic girdle length | 0.951 | 0.952 | 1.000 | 0.932 | 0.88 | 0.822 | 0.826 | 0.801 | 0.671 | 0.443 |
| Ascending process length | 0.931 | 0.945 | 0.934 | 1.000 | 0.922 | 0.795 | 0.796 | 0.788 | 0.653 | 0.393 |
| Ascending process width | 0.907 | 0.897 | 0.889 | 0.930 | 1.000 | 0.766 | 0.783 | 0.757 | 0.699 | 0.478 |
| 2nd dorsal spine | 0.838 | 0.894 | 0.853 | 0.823 | 0.793 | 1.000 | 0.81 | 0.96 | 0.742 | 0.638 |
| Mean lateral plate length | 0.881 | 0.840 | 0.853 | 0.816 | 0.797 | 0.853 | 1.000 | 0.785 | 0.872 | 0.696 |
| 1st dorsal spine | 0.818 | 0.887 | 0.836 | 0.820 | 0.784 | 0.967 | 0.83 | 1.000 | 0.725 | 0.605 |
| Body depth | 0.782 | 0.698 | 0.736 | 0.700 | 0.722 | 0.806 | 0.905 | 0.784 | 1.000 | 0.889 |
| Standard length | 0.621 | 0.534 | 0.579 | 0.514 | 0.559 | 0.736 | 0.788 | 0.700 | 0.933 | 1.000 |

MANOVA suggests that the types vary in at least one of the traits (Pillai’s Trace omnibus F_4, 856_ = 17.58. p<< 0.001; Fig. 4). All antipredator armor traits vary among the types (one-way ANOVA for each trait; Table 2; Fig. 4) and trait value tend to be smallest for stickleback from the Narrows Pond type for all traits except standard length. Fig.Fig.Fig.

**Figure 4.**
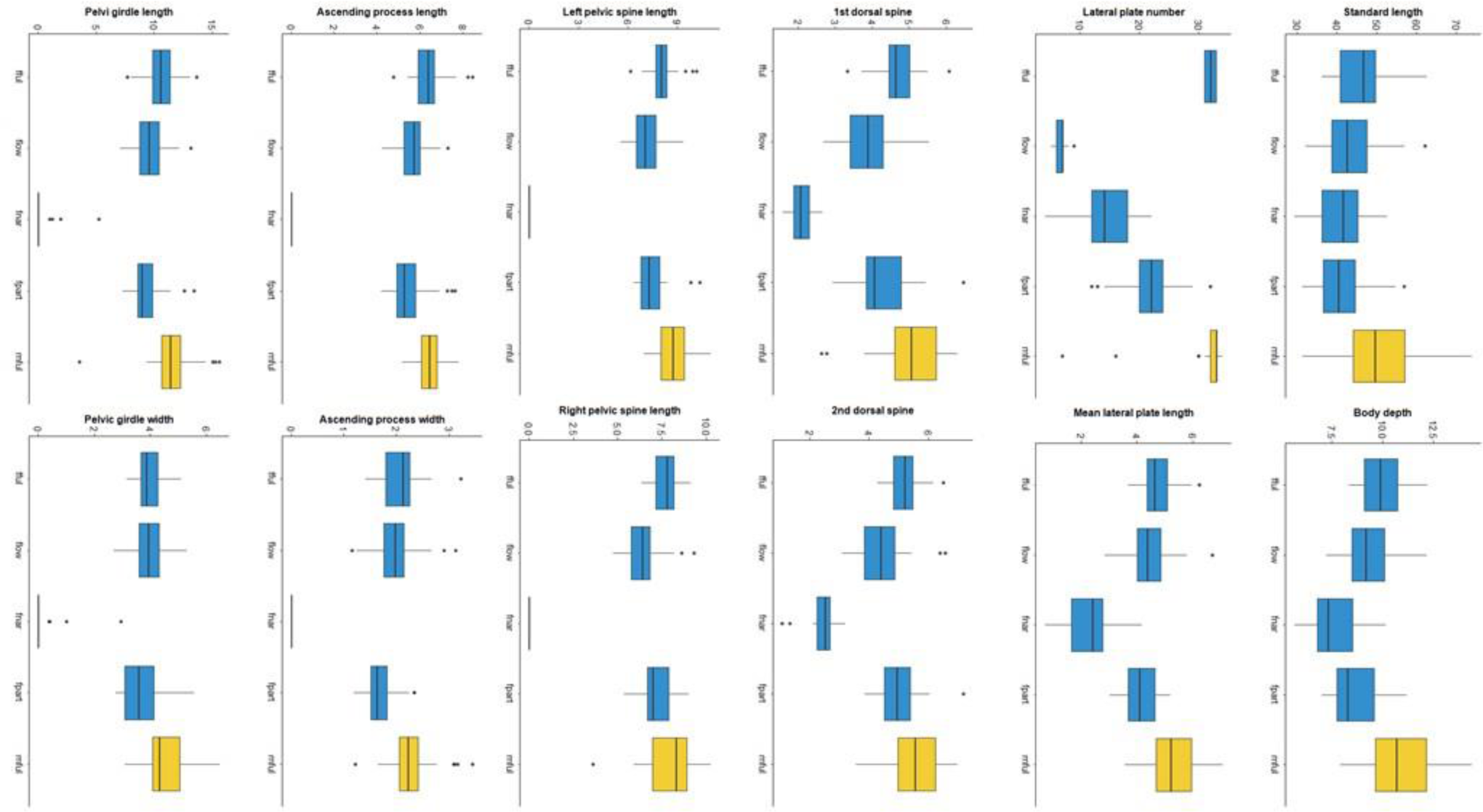
Box plots showing median (line in box), Q1 and Q3 (bottom and top hinges respectively), and extremes (1.5 times the lower and upper hinges) of variables measured for each stickleback type sampled. Yellow symbols indicates marine locations and blue symbols indicate freshwater locations.

**Figure 5.**
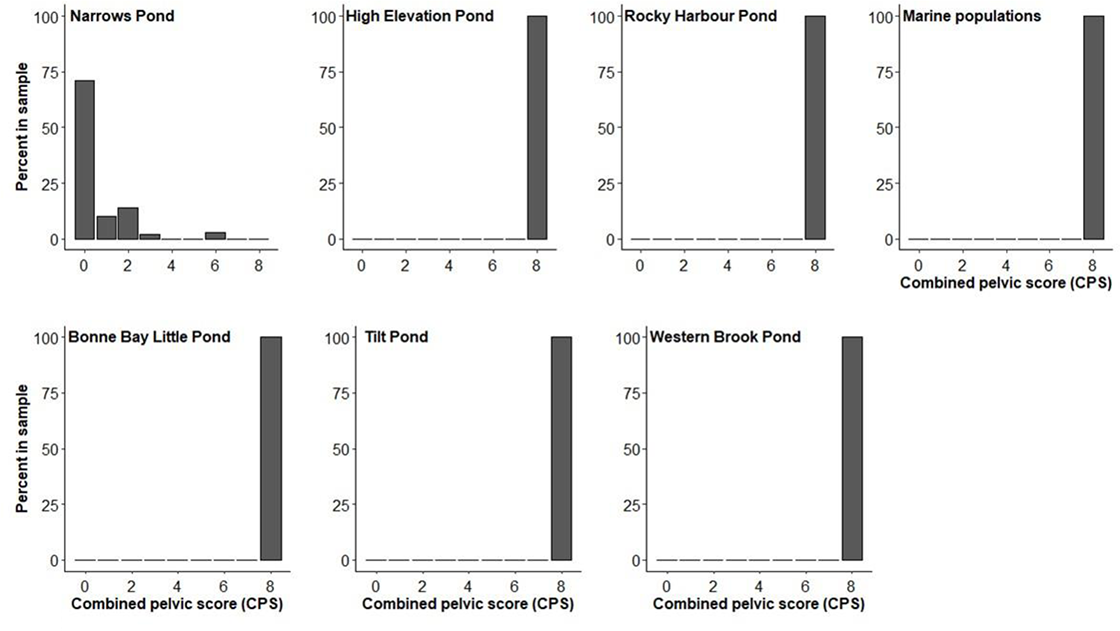
Percent occurrence of combined pelvic score (CPS) for each population sampled. CPS was is based on the criteria described in Klepaker and Østbye (2008).

**Table 2.**
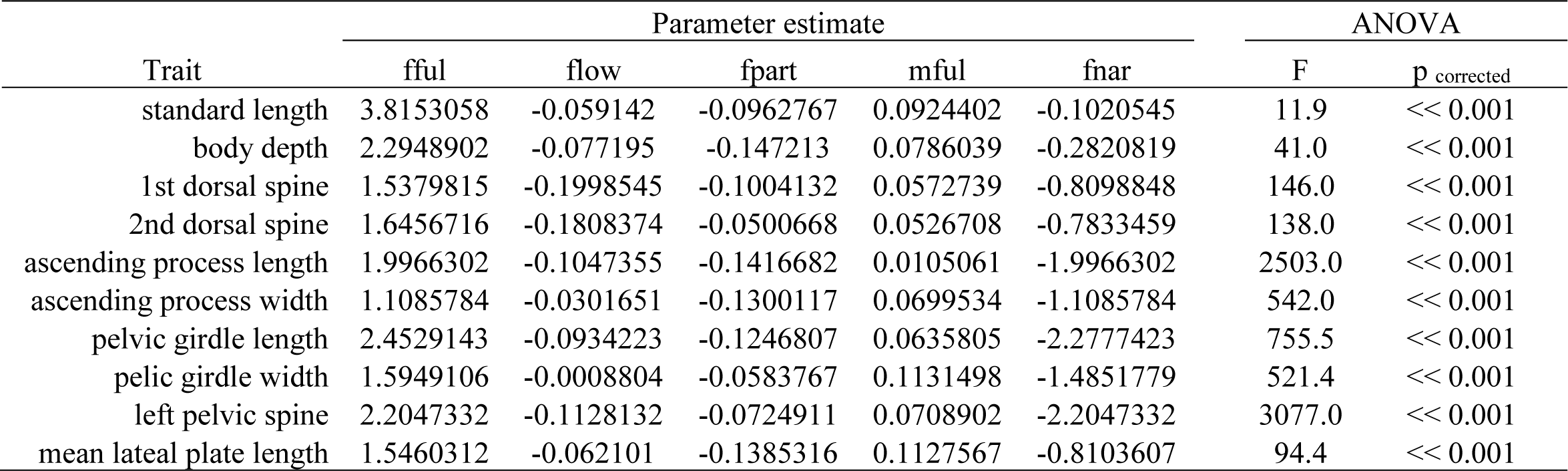
Effect size (relative to fful type) and results of one-way ANOVA (F and Bonferoni corrected p-value) comparing ln-transformed trait values among the five stickleback types in this study.

The combined pelvic score for each of the five groups is summarized in Fig. 6. Nearly 75% of the individuals assessed from Narrows Pond had no expression of any pelvic girdle components. Most of the remaining roughly 25% of the individuals showed little pelvic girdle expression (CPS = 1 to 2), with only a few individuals scoring as high as CPS = 6). All fish from the other populations showed completed pelvic girdle expression (CPS = 8).

## Discussion

We compared the armor traits of Narrows Pond stickleback to several other nearby stickleback populations. Stickleback in Narrows Pond have by and large lost their entire pelvic girdle and have overall reduced anti-predator armor traits. Pelvic girdle loss and armor reduction is rare among threespine stickleback populations, but has been observed in in populations in western North America (British Columbia: Reimchen 1984, Alaska: Bell and Orti 1994), Europe (Scotland: Giles 1983; Campbell 1984, Coyle et al. 2007; Norway: Klepaker and Ostybe 2008, Klepaker et al. 2012; and eastern North America: Edge and Coad 1983, this study). This is the first population with pelvic girdle loss/reduction in Newfoundland and only the third in eastern North America, with the other two being within 4 km of each other.

Surveys that have examined pelvic reduction in threespine stickleback show variability in the extent of pelvic reduction in the populations where reduction occurs. For example, 97% of individuals sampled in Serendipity Lake, BC possessed no pelvic spines whereas only 65% of individuals in Rouge Lake, B.C. had no pelvic spines (Reimchen 1984). The proportion of individuals with pelvic reduction in each of the four Norwegian populations in Klepaker and Østbye (2008) ranged from 4.7% to 68% of all individuals sampled in each lake. The Scottish populations with pelvic reduction also appear to be mixed with individuals that have a complete pelvic girdle, though it is hard to estimate proportions based on the way the data was presented in those studies (Giles 1983; Campbell 1984). We scored 100 individuals from Narrows Pond and did not find a single individual with a complete pelvic girdle. We did observe 3 individuals with CPS = 6. However, we expect that how we scored may have been a little more liberal than other studies and likely our scoring is a little high especially in comparison to that of Klepaker et al. (2012). Regardless, it is clear that the Narrows Pond population demonstrates extreme pelvic reduction, and is certainly the most extreme case of pelvic reduction in the eastern North American stickleback lineages.

Two factors have been hypothesized to promote armor and pelvic girdle reduction, low environmental calcium availability and predator absence (e.g. Bell et al. 1993, Lescak et al. 2012; Reimchen et al. 2013). However, neither of these explanations is likely for the Narrow’s pond population. Brook char (*Salvelinus fontinalis*), a likely predator of threespine stickleback, is abundant in Narrows Pond. One of us (R. Scott) has observed schools of brook char numerous times while snorkeling and a 15 minute gill net set led to the capture of two brook char that were approximately 25 cm long. Also, the surficial geology surrounding Narrows Pond is dominated by dolostone (DeGrace 1974) which, when dissolved leads to elevated calcium availability in aquatic systems (e.g. Hinder et al. 2003). Lac Croche and Lac Rond—two lakes within Québec’s Parc national du Lac-Témiscouata, and the only other documented populations of pelvic-reduced threespine stickleback in eastern N. America—also sit in a region of high-calcium geology (the Lac Croche and Saint-Léon formations of the Gaspé Belt) meaning calcium is very unlikely to be limiting (Ministére du Développement durable, Environnement et Parcs 2008). However, the frequency of pelvic reduction in Lac Rond fell dramatically in the decades following the introduction of brook trout (Edge and Coad 1983, LaCasse and Aubin-Horth 2012). This makes the Narrows Pond population unique in its sustained pelvic reduction despite the presence of a predator and abundantly available calcium.

Unlike lateral plate morphs, for which the low-plate alleles are typically maintained at low frequencies in marine populations and transported between freshwater drainages via the marine environment (Roberts Kingman et al. 2021), pelvic reduction typically induced by *de novo* mutations in a particularly fragile region of the genome (Xie et al. 2019). Irrespective of the source of the mutation regulating this dramatic morphological divergence in the Narrows Pond population, though, the source of selection maintaining it under conditions that would typically select against reduced pelvises remains obscure.

Surveys of stickleback armor show a broad range of variability among freshwater populations. Scott et al. (2022) found variability among 52 threespine stickleback populations from western Newfoundland that parallels variability observed other parts of the species’ range; marine populations, as elsewhere, did not vary across the sample range and freshwater populations varied both in armor expression and body shape. We used stickleback from eight populations from Scott et al. (2022) representing marine and freshwater locations. The freshwater locations chosen included populations that are monomorphic low, partial, or complete with regard to lateral armor plate characteristics. While typical in the rest of the world, this variation in plate morphs appears higher in Newfoundland than in the rest of eastern Canada (Hagen and Moodie 1982). Narrows Pond stickleback are clearly different in all aspects of armor expression relative to the reference populations; Narrows Pond stickleback have a reduced pelvis (completely lost in most individuals sampled), and have smaller spines and lateral plates, and they have shallower bodies.

## Acknowledgements

J. Moriera, M. Graham, D. Budgill, and W. Rauch-Davis assisted with collections and W. Rauch Davis also assisted with image capture and digitization. This research was supported by the Grenfell Campus Research Fund and NSERC USRA (to C.A. Trask), approved by the Memorial University Animal Care committee, and conducted under collection permits issued by Parks Canada, and Department of Fisheries and Oceans Canada.

## Data availability

Data have been uploaded to Borealis and can be accessed at doi:10.5683/SP3/CT1SRW following publication of this manuscript in a per-reviewed journal.

## Ethics declaration

All field collections were carried out under Memorial University Animal Care approval (file numbers 20090160 and 2018015) and under Scientific Collection permits issued by Parks Canada and department of Fisheries and Oceans Canada.

## References

Arostegui MC, Quinn TP (2019) Reliance on lakes by salmon, trout and charr (Oncorhynchus, Salmo, and Salvelinus): An evaluation of spawning habitats, rearing strategies and trophic polymorphisms. Fish and Fisheries 20:775–794 doi:10.1111/faf.12377

Barrett RDH (2010) Adaptive evolution of lateral plates in three-spined stickleback Gasterosteus aculeatus: a case study in functional analysis of natural variation. Journal of Fish Biology 77(2):311–328 doi:10.1111/j.1095-8649.2010.02640.x

Bell MA (1987) Interacting evolutionary constraints in pelvic reduction of threespine sticklebacks, Gasterosteus aculeatus (Pisces, Gasterosteidae). Biological Journal of the Linnean Society 31:347–382 doi: 10.1111/j.1095-8312.1987.tb01998.x

Bell MA, Orti G (1994) Pelvic reduction in threespine stickleback from cook inlet lakes, geographical distribution and intrapopulation variation. Copeia(2):314–325

Bell MA, Orti G, Walker JA, Koenings JP (1993) Evolution of pelvic reduction in threespine stickleback fish, a test of competing hypotheses. Evolution 47(3):906–914 doi:10.2307/2410193

Campbell RN (1985) Morphological variation in the 3-spined stickleback (Gasterosteus aculeatus) in Scotland. Behaviour 93:161–168 doi:10.1163/156853986x00838

Coyle SM, Huntingford FA, Peichel CL (2007) Parallel evolution of Pitx1 underlies pelvic reduction in Scottish threespine stickleback (Gasterosteus aculeatus). Journal of Heredity 98(6):581–586 doi:10.1093/jhered/esm066E

DeGrace, JR (1974) Limestone resources of Newfoundland and Labrador, Report 74-2. Newfoundland Department of Mines and Energy, Mineral Development Division.

Edge TA, Coad BW (1983) Reduction of the pelvic skeleton in the threespine stickleback, Gasterosteus aculeatus, in 2 lakes of Québec. Canadian Field-Naturalist 97(3):334–336

Fang B, Kemppainen P, Momigliano P, Feng X, Merilä J (2020) On the causes of geographically heterogeneous parallel evolution in sticklebacks. Nature Ecology & Evolution 4:1105–1115 doi:10.1038/s41559-020-1222-6

Foster SA, Scott RJ, Cresko WA (1998) Nested biological variation and speciation. Philosophical Transactions of the Royal Society B-Biological Sciences 353(1366):207–218 doi:10.1098/rstb.1998.0203

Giles N (1983) The possible role of environmental calcium levels during the evolution of phenotypic diversity in Outer Hebridean populations of the 3-spined stickleback, Gasterosteus aculeatus. Journal of Zoology 199(4):535–544 doi: 10.1111/j.1469-7998.1983.tb05104.x

Hagen DW and Moodie GEE (1982) Polymorphism for plate morphs in Gasterosteus aculeatus on the east coast of Canada and an hypothesis for their global distribution. Canadian Journal of Zoology 60:1032–1042 doi: 10.1139/z82-144

Hindar A, Nilsen P, Skiple A, Hogberget R (1995) Counteractions against acidification in forests ecosystems, effects on stream water quality after dolomite application to forest soil in Gjerstad, Norway. Water Air and Soil Pollution 85(2):1027–1032 doi:10.1007/bf00476965

Horst AM, Hill AP, Gorman KB (2020). palmerpenguins: Palmer Archipelago (Antarctica) penguin data. R package version 0.1.0, https://allisonhorst.github.io/palmerpenguins/. doi: 10.5281/zenodo.3960218.

Roberts Kingman GA, Le D, Jones FC, Desmet D, Bell MA, Kingsley DM (2021) Longer or shorter spines: Reciprocal trait evolution in stickleback via triallelic regulatory changes in Stanniocalcin2a. PNAS 118: e2100694118 doi: 10.1073/pnas.2100694118.

Klepaker T, Østbye K, Bell MA (2013) Regressive evolution of the pelvic complex in stickleback fishes: a study of convergent evolution. Evolutionary Ecology Research 15:413–435.

Klepaker T, Østbye K, Bernatchez L, Vollestad LA (2012) Spatio-temporal patterns in pelvic reduction in threespine stickleback (Gasterosteus aculeatus L.) in Lake Storvatnet. Evolutionary Ecology Research 14(2):169–191

Klepaker TO, Østbye K (2008) Pelvic anti-predator armour reduction in Norwegian populations of the threespine stickleback: a rare phenomenon with adaptive implications? Journal of Zoology 276(1):81–88 doi:10.1111/j.1469-7998.2008.00471.x

LaCasse J, Aubin-Horth N (2012) A test of the coupling of predator defense morphology and behavior variation in two threespine stickleback populations. Current Zoology 58(1):53–65 doi:10.1093/czoolo/58.1.53

Lescak EA, von Hippel FA, Lohman BK, Sherbick ML (2012) Predation of threespine stickleback by dragonfly naiads. Ecology of Freshwater Fish 21(4):581–587 doi:10.1111/j.1600-0633.2012.00579.x

Lleornart J, Salat J, Torres GJ (2000) Removing allometric effects of body size in morphological analysis. J. Theor. Biol. 205: 85–93.

Losos JB (2010) Adaptive Radiation, Ecological Opportunity, and Evolutionary Determinism. American Naturalist 175(6):623–639 doi:10.1086/652433

Ministére du Développement durable, Environnement et Parcs (2008) Parc national du Lac-Témiscouata: États des connaissance. ISBN : 978-2-550-52191-4

Quinn, G. P., & Keough, M. J. (2002). Experimental design and data analysis for biologists. Cambridge, UK: Cambridge University Press.

Reimchen TE (1983) Structural relationships between spines and lateral plates in threespine stickleback (Gasterosteus aculeatus). Evolution 37(5):931–946 doi:10.2307/2408408

Reimchen TE (1984) Status of unarmored and spine-deficient populations (Charlotte unarmored stickleback) of threespine stickleback, Gasterosteus sp, on the Queen Charolotte Islands, British Columbia. Canadian Field-Naturalist 98(1):120–126

Reimchen TE, Bergstrom C, Nosil P (2013) Natural selection and the adaptive radiation of Haida Gwaii stickleback. Evolutionary Ecology Research 15(3):241–269

Robinson BW, Parsons KJ (2002) Changing times, spaces, and faces: tests and implications of adaptive morphological plasticity in the fishes of northern postglacial lakes. Canadian Journal of Fisheries and Aquatic Sciences 59(11):1819–1833 doi:10.1139/f02-144

Robinson BW, Wilson DS (1994) Character release and displacement in fishes - a neglected literature. American Naturalist 144(4):596–627 doi:10.1086/285696

R Core Team (2022) R: A language environment for statistical computing. R Foundation for Statistical Computing, Vienna, Austria, URL: https://www.R-project.org/.

RStudio Team (2020) RStudio: integrated development for R. RStudio, PBS, Boston, MA. URL: http://www.rsctudio.com.

Schneider C, Rasband W, Eliceiri K (2012) NIH Image to Image J: 25 years of image analysis. Nat. Methods 9: 671–675. doi: http://doi.org/10.1038/nmeth.2089.

Scott, RJ, Haines, GE, Biedak, N, Baker, JA (2022) Variation in morphology among populations of threespine stickleback (Gasterosteus aculeatus) from western Newfoundland, Canada. Submitted to Environmental Biology of Fishes

Song JK, Parenti LR (1995) Clearing and staining whole fish specimens for simultaneous demonstration of bone, cartilage, and nerves. Copeia(1):114–118

Stuart YE, Veen T, Weber JN, Hanson D, Ravinet M, Lohman BK, Thompson CJ, Tasneem T, Doggett A, Izen R, Ahmed N, Barrett RDH, Hendry AP, Peichel CL, Bolnick DI (2017) Contrasting effects of environment and genetics generate a continuum of parallel evolution. Nature Ecology & Evolution 1:0158 doi:10.1038/s41559-017-0158

Xie TK, Wang G, Thompson AC, Wucherpfennig JI, Reimchen TE, MacColl ADC, Schluter D, Bell MA, Vasquez KM, Kingsley DM (2019) DNA fragility in the parallel evolution of pelvic reduction in stickleback fish. Science 363(6422):81–84 doi: 10.1126/science.aan1425

